# Microbial ecology of Atlantic salmon, *Salmo salar*, hatcheries: impacts of the built environment on fish mucosal microbiota

**DOI:** 10.1101/828749

**Authors:** Jeremiah J Minich, Khattapan Jantawongsri, Colin Johnston, Kate Bowie, John Bowman, Rob Knight, Barbara Nowak, Eric Allen

**Affiliations:** Marine Biology Research Division, Scripps Institution of Oceanography; Institute for Marine and Antarctic studies, University of Tasmania; Tassal Operations Pty Ltd. Hobart, Tasmania; Tasmanian Institute of Agriculture, University of Tasmania; Center for Microbiome Innovation, University of California San Diego; Department of Pediatrics, University of California San Diego

## Abstract

Successful rearing of fish in hatcheries is critical for conservation, recreational fishing, and commercial fishing through wild stock enhancements, and aquaculture production. Flow through (FT) hatcheries require more water than Recirculating-Aquaculture-Systems (RAS) which enable up to 99% of water to be recycled thus significantly reducing environmental impacts. Here, we evaluated the biological and physical microbiome interactions of the built environment of a hatchery from three Atl salmon hatcheries (RAS n=2, FT n=1). Six juvenile fish were sampled from tanks in each of the hatcheries for a total of 60 fish across 10 tanks. Water and tank side biofilm samples were collected from each of the tanks along with three salmon body sites (gill, skin, and digesta) to assess mucosal microbiota using 16S rRNA sequencing. The water and tank biofilm had more microbial richness than fish mucus while skin and digesta from RAS fish had 2× the richness of FT fish. Body sites each had unique microbial communities (P<0.001) and were influenced by the various hatchery systems (P<0.001) with RAS systems more similar. Water and especially tank biofilm richness was positively correlated with skin and digesta richness. Strikingly, the gill, skin and digesta communities were more similar to the origin tank biofilm vs. all other experimental tanks suggesting that the tank biofilm has a direct influence on fish-associated microbial communities. The results from this study provide evidence for a link between the tank microbiome and the fish microbiome with the skin microbiome as an important intermediate.

**IMPORTANCE:** Atlantic salmon, *Salmo salar*, is the most farmed marine fish worldwide with an annual production of 2,248 million metric tonnes in 2016. Salmon hatcheries are increasingly changing from flow through towards RAS design to accommodate more control over production along with improved environmental sustainability due to lower impacts on water consumption. To date, microbiome studies on hatcheries have focused either on the fish mucosal microbiota or the built environment microbiota, but have not combined the two to understand interactions. Our study evaluates how water and tank biofilm microbiota influences fish microbiota across three mucosal environments (gill, skin, and digesta). Results from this study highlight how the built environment is a unique source of microbes to colonize fish mucus and furthermore how this can influence the fish health. Further studies can use this knowledge to engineer built environments to modulate fish microbiota for a beneficial phenotype.

## INTRODUCTION

Aquaculture is the fastest growing agricultural industry now producing over 50 % of seafood by volume globally (1). While freshwater systems currently outproduce marine systems (51.4 MT vs. 28.7 MT; 2016), marine aquaculture has a tremendous potential to expand with estimates of theoretical production of 15 billion tonnes (522 × increase) (2, 3). One of the primary challenges to scaling aquaculture production is improving seed quality by increasing survival rates and strengthening immune development of larvae and juveniles in the hatchery environment (4). This becomes challenging particularly when there are an estimated 369 different species of fish currently grown for commercial aquaculture with additional species in experimental production (3). In terms of global aquaculture production, Atlantic salmon, *Salmo salar*, ranks first among marine fish and is the 9^th^ largest aquaculture fish species overall (3). Global growth in Atlantic salmon production has primarily been driven by technological advancements in automated feeding machines, reduced reliance on fishmeal based feeds, selective breeding to reduce growout time to market from three years to one year (at sea), and in the Northern hemisphere, disease control through commercial adoption of vaccine development along with biological control of parasite infection using cleaner fish (5). Note that neither Australia nor New Zealand has issues with sea lice, but like the Northern hemisphere does have amoebic gill disease. Improvements in hatchery technology is further reducing the environmental footprint of aquaculture. Optimizing hatchery conditions is also important for mariculture, capture fisheries, recreational fisheries, and conservation as many government programs rely on ocean enhancement efforts to replenish wild populations. For example, in 2018 29 Alaska salmon hatcheries used for ocean enhancement contributed to 34% of commercial harvest worth 453 million USD (6). Understanding the factors for which hatchery reared salmon exhibit altered performance compared to wild salmon including faster growth rates, lower age to maturity, higher overall survival, lowered lifetime reproductive success, and increased aggression including competitiveness, may be important for improved ocean enhancement. (7–9).

Salmon are reared in two primary types of freshwater hatchery systems: flow through (FT) which requires continuous new water and recirculating aquaculture systems (RAS) where up to 99% of water is recycled. Flow through (FT) hatcheries however, take in and release relatively large volumes of water, usually from natural surface waters, and require water treatment and settlement systems. The RAS systems have the potential to significantly reduce freshwater requirements and thereby lower environmental impacts. One major concern for RAS systems over traditional flow-through (FT) systems is a potential impact on fish health which is in part thought to be due to microbial dysbiosis either to the fish or the environment (water). Atlantic salmon hatcheries are primarily built near freshwater inputs such as streams or rivers whereby water is filtered and either continually flowed through the tanks at approximately 300% daily or used to replenish the RAS tanks at 2-7% daily. During the freshwater stage in both hatchery systems, juvenile salmon, parr, are reared in circular tanks ranging from 3 m to 10 m in diameter and 2-5 m in depth made from fiberglass, concrete, or other materials equipped with Oxygen injectors (aerators). This period is crucial for salmon survival as disease outbreaks can cause costly die offs in the system. Compared to flow through systems, enclosed RAS systems have the benefit of requiring 93% less water from the environment and a 26-38% reduced eutrophication on the environment (10, 11), but can also be more costly in energy use (24-40% higher) (12). Because RAS systems are enclosed batch systems, biosecurity is theoretically improved as conditions can be regulated and controlled much easier than in FT systems. In addition, the feed conversion ratio (FCR) can be lower in a RAS system due to ability to control all variables such as temperature and salinity (10). RAS systems may enable establishment of stable, slow growing, bacterial communities in hatchery systems which can improve survival rates in cod (13). Other studies however have suggested that water quality (higher recirculating microbial loads, accumulation of metabolites, or accumulation of heavy metals from feeds) in RAS systems was detrimental for larvae survival and/or growth of common carp (14), sea bass (15), and nile tilapia (16). For post-smolt Atlantic salmon reared in a RAS system, both salinity (12, 22, and 32 ppt) and time (3, 4.5, 7 months) influenced microbial communities of the water column, while the tank biofilm (which differed from water column) remained stable (17). Since microbial communities are indicated as an important factor in RAS water quality, and thereby fish health, it is important to understand how microbiomes of both the built environment along with the fish mucus are influenced in RAS versus FT systems.

The importance of mucus (gill, skin, GI) microbiome (collection of microbial eukaryotes, bacteria, archaea, and viruses) to animal health has been well documented and it is through mechanisms such as competitive exclusion, production of antimicrobial compounds, and microenvironment control to reduce pathogen growth and colonization (18). Mucosal environments including the gill (19), skin (20), and gut (21) serve as important physical barriers for disease and are important part of immune response. The skin and gut microbiomes of Atlantic salmon are unique, differ by life stage (parr, smolt, adult), and differ depending upon rearing environment (wild vs. hatchery). Furthermore, water has been shown to primarily influence the skin community (22) which is further exemplified during migration from freshwater to saltwater (23). Gut microbiomes of Atlantic salmon is primarily driven by the life stage rather than environment (24, 25) in the wild which has been hypothesized to be due to changes in diet along with increased consumption of water during the marine stage (26). The hatchery built environment is a unique microbial habitat which has largely remained unexplored (27, 28). Understanding the relationship between the built environment of the hatchery along with the mucosal microbiome of the fish may be important for predicting fish health.

The purpose of this experiment was to evaluate how hatchery type (FT vs RAS) influences the microbial community of fish mucus and subsequently fish health. This is the first study to holistically evaluate the gill, skin, and gut microbiome of Atlantic salmon. We further combined histological analyses of mucosal sites to connect microbial changes to mucus health.

## METHODS

Six fish were randomly sampled from each of 10 tanks across three freshwater hatcheries. Fish were collected and euthanized using the AQUI-S by husbandry technicians according to label instructions. Biometrics including total length, mass, and condition factor were measured. Tank conditions such as water temperature, salinity, and diameter were recorded and can be found in the metadata file.

### Histology

Samples of gills, skin and intestine, fixed in 10% buffered formalin were trimmed, processed using standard protocols for histology and embedded in paraffin. Sections of 4 μm were cut and one section was used for each individual fish. The sections were stained with Alcian Blue/ Periodic Acid – Schiff (AB/PAS) at pH 2.5 to quantify mucous cells in the gills per inter-lamellar unit (ILU) under a bright field light microscope (Leica DM1000, Hamburg, Germany) (29, 30), to count the number of mucous cells in the skin and the number of intestinal mucous cells, normalized per area (31). For the gut morphometric measurements of the fold height, mucosa thickness, fold width, muscularis thickness, fold height, mucosa thickness, fold width, and muscularis thickness were done as previously described (32). Intestine sections stained with haematoxylin and eosin (H&E) were analysed using Image-Pro Premier software. Ten intestinal folds from one section from each region were included in the analysis.

### Microbiome processing

For each of the ten tanks, six fish were collected individually using hand nets and placed directly into a sterile sampling bucket and anesthetized using AQUI-S. The mucosal microbiome was sampled as follows: gill by swabbing the second gill arch on the left lateral side; skin by swabbing a 2 cm × 2 cm area posterior of the operculum on the left lateral side under the dorsal fin; and digesta by massaging the GI until a fecal pellet emerged. Swabs were then placed directly into a 2 ml PowerSoil tube and frozen at −20 °C. In addition to biological samples, two environmental samples were taken per tank including a 2 cm × 2 cm swab of the inside of the tank just below the water line (biofilm) along with a 400 ul bulk water sample from the tank. In total three body sites across 60 fish (180 samples) along with two environmental samples across ten tanks (20 samples) were collected and processed for microbiome analysis. In addition, 21 technical controls were included.

DNA extraction was performed at University of Tasmania Hobart using the EarthMicrobiomeProject protocols (earthmicrobiome.org), specifically using the ‘manual single-tube’ MoBio PowerSoil kit as to reduce well-to-well contamination (33). A total of 21 positive controls of a microbial isolate (replicates of 10 fold serial dilutions) were processed alongside the samples and then used to determine sample success rate by calculating the sample exclusion criteria based on read counts described in the Katharoseq method (27). Samples were processed in triplicate 5 ul PCR reactions (34)[PMID: 30417111] using the 16S v4 515/806 primers (35, 36) and then pooled at equal volume according to Katharoseq (27). The final amplicon pool was processed using the Qiagen PCR cleanup kit following EMP protocols and sequenced on a MiSeq 2×250 bp run (37). Sequencing runs were processed in Qiita (38) using Qiime2 commands (39). Samples were trimmed to 150 bp and then processed through the deblur (40) pipeline which generates unique, single ASVs (amplicon sequence variants). To determine which samples had been sequenced successfully, the Katharoseq method (27), developed for low biomass sequencing, was applied. The cutoff value for composition of a sample aligning to the target within the positive controls was 90%. In this case, the cutoff value was 405 reads, but we rarified to 1000 reads to have higher depth of sequencing. Within Qiita, samples which did not have histology metadata were excluded.

### Statistical analysis

Alpha diversity was calculated using richness (total observed unique ASVs) and Faith’s Phylogenetic Diversity. Differences between body sites and environmental variables was tested using non-parametric Kruskal-Wallis test with the Benjamini Hochberg FDR 0.05 (41, 42). Correlations between richness of environmental variables (tank and bioiflm) and salmon body site was calculated using linear regression. For beta diversity we used weighted and unweighted UniFrac (43, 44). Multivariate statistical testing of both continuous and categorical variables was performed using ADONIS within Qiime(45). Pairwise statistical comparisons of beta diversity measures were calculated using Mann-Whitney while multiple comparisons conducted using Kruskal-Wallis test. To identify correlations between histological measures and specific microbes, a non-parametric, Spearman correlation was calculated for both the entire dataset using the Calour analysis tool (46).

## RESULTS

A total of 60 fish were sampled from three unique hatcheries, one flow through (FT) and two recirculating aquaculture systems (RAS). Within the hatcheries, a total of six fish were sampled from each of ten unique tanks. To evaluate health status, fish were examined for histopathological measurements within the gill, skin, and gastrointestinal tract. In addition, the mucosal microbiome of three body sites (gill, skin, and digesta) was sampled across all 60 fish along with environmental controls including the tank water and tank-associated biofilms. After calculating sample cutoff measures and rarefying to 1000 reads, a total of 185 samples passed QA/QC resulting in a total of 6,197 total unique ASVs (Supp Figure S1). Failures were not associated with any particular hatchery system (success rate: 72/78 RAS 1, 56/60 RAS 2, 56/60 FT) or body site (success rate: 56/60 gill, 58/60 skin, 55/60 digesta). A total of 37 microbial Phyla were represented in the dataset including one archaea (Euryarchaeota) and one eukaryote (Apicomplexa) (Supp Figure S2). Digesta samples generally had higher levels of Cyanobacteria, Firmicutes, Actinobacteria, and Fusobacteria whereas the skin and gill were enriched with Bacteroidetes, Verrucomicrobia, and Acidobacteria. Across all body sites and built environment samples, Proteobacteria was most dominant.

Statistical analyses of community composition revealed that body sites along with hatchery system and further tank replicates were all significant drivers of community composition with body site (P=0.001, R2 = 0.127 Unweighted Unifrac; P=0.001, R2 = 0.340 Weighted UniFrac) being the strongest (Table 1). Furthermore, when stratifying for each body site (gill, skin, and gut), microbial communities were significantly influenced by both hatchery location and across individual tanks using both Unweighted and Weighted UniFrac (Table 1).

**Table 1.** Multivariate statistical testing of drivers of microbial beta diversity

|  |  |  |  |  |  |  |  |  |  |  |  |
| --- | --- | --- | --- | --- | --- | --- | --- | --- | --- | --- | --- |
| Unweighted UniFrac | metadata column | Variable_type | method | Combined |  | gill |  | skin |  | gut |  |
|  |  |  |  | R2 | P | R2 | P | R2 | P | R2 | P |
| Body site | sample_type | categorical | adonis | 0.127 | **0.001** | Nan | Nan | Nan | Nan | Nan | Nan |
| Hatchery system | ylk_tank_system | categorical | adonis | 0.062 | **0.001** | 0.124 | **0.001** | 0.172 | **0.001** | 0.091 | **0.001** |
| Tank number | ylk_sbt_tank_number | categorical | adonis | 0.110 | **0.001** | 0.268 | **0.001** | 0.294 | **0.001** | 0.284 | **0.001** |
| Weighted UniFrac |  |  |  |  |  |  |  |  |  |  |  |
| Body site | sample_type | categorical | adonis | 0.340 | **0.001** | Nan | Nan | Nan | Nan | Nan | Nan |
| Hatchery system | ylk_tank_system | categorical | adonis | 0.053 | **0.001** | 0.231 | **0.001** | 0.229 | **0.001** | 0.084 | **0.031** |
| Tank number | ylk_sbt_tank_number | categorical | adonis | 0.119 | **0.002** | 0.423 | **0.001** | 0.388 | **0.001** | 0.433 | **0.001** |

Microbial diversity differs according to sample type with water samples having the highest richness (P=0.0015, KW 17.52) and phylogenetic diversity (P=0.0021, KW 16.79) (Figure 1a-b). When comparing only fish mucus samples, the gill had less richness than the skin and digesta (P=0.0056, KW 10.37) and lower phylogenetic diversity than the skin (P=0.0279, KW 7.16). Microbial composition as assessed using Unweighted UniFrac distances, was primarily driven by sample type followed by hatchery system with samples from the RAS generally being more similar than the FT hatchery (Table 1, Figure 1c-d). In addition, water and biofilm samples were highly distinguishable between the hatchery systems, particularly RAS vs. FT and clustered more closely to gill and skin samples indicating that gill and skin microbiomes were more closely related to the built environment.

**Figure 1.**
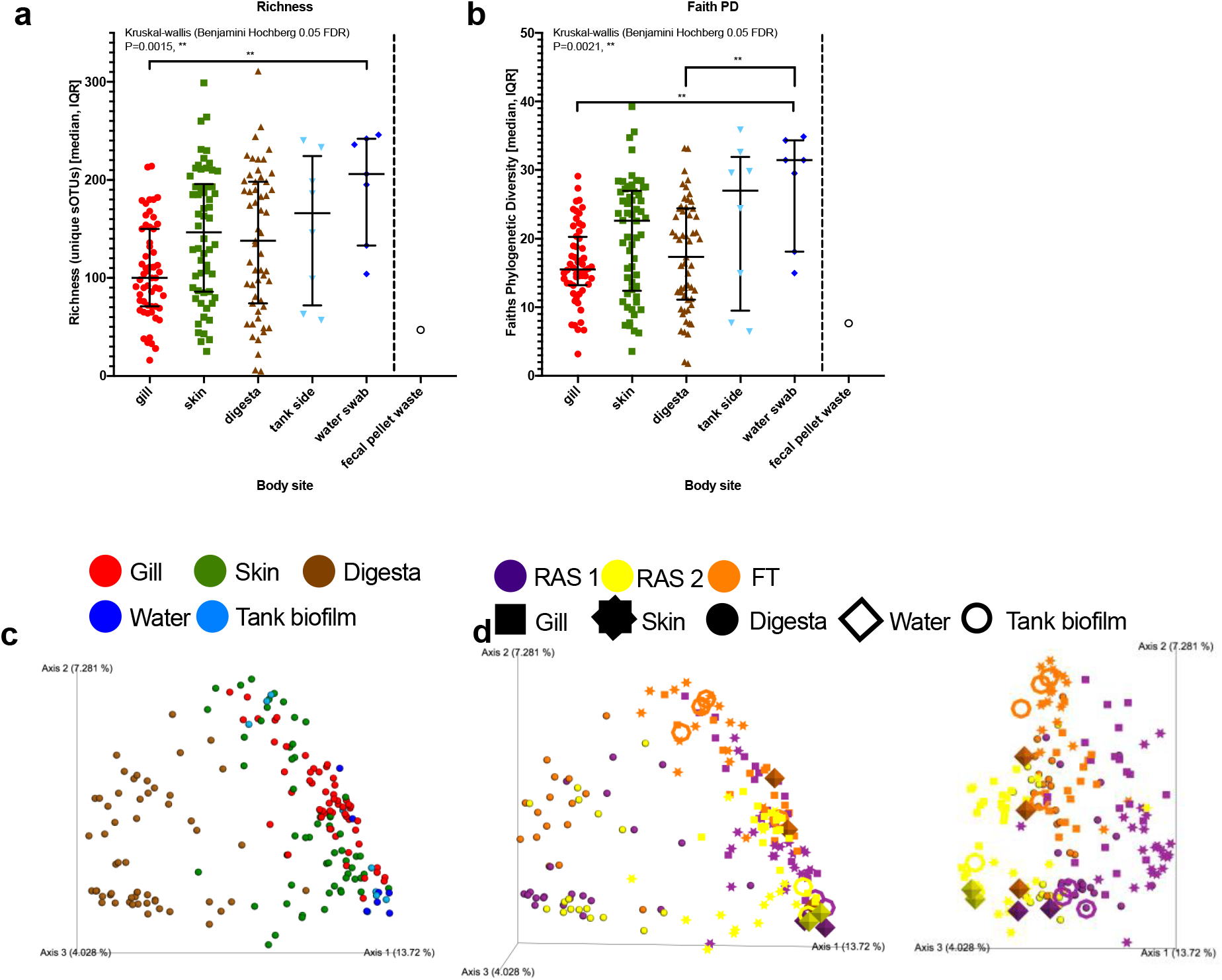
Microbial ecology (16S rRNA) of three tanks each from three hatchery systems (water and tank biofilm) and Atlantic salmon gill mucus, skin mucus, and digesta. Alpha diversity measures of a) total richness and b) Faiths Phylogenetic Diversity evaluated by non-parametric Kruskal-Wallis test with Benjamini-Hochberg 0.05 FDR. Beta diversity measures of unweighted UniFrac distances colored by c) sample type and by d) hatchery system.

We next assessed how facility type influenced the microbiome of both the fish body sites and the built environment. Microbial richness of skin, digesta, tank biofilm and tank water was generally higher in the RAS systems compared to FT (Figure 2a). When comparing only fish body sites, both skin (P<0.0001, KW 21.16) and digesta (P=0.0058, KW 10.29) richness was significantly different across hatcheries with RAS systems having approximately 2× more sOTUs associated with skin and digesta compared to FT (Figure 2b). Post-hoc multiple comparison tests demonstrated that for skin both RAS1 and RAS2 richness was greater than FT whereas for digesta only RAS1 was higher than FT (Figure 2b). Compositionally, the microbial communities were significantly different across hatcheries for all samples combined (Table 1). When only analyzing microbial communities of specific body sites like gill, skin, and digesta, a hatchery specific microbiome was still observed (Figure 2c-e). The hatchery specific microbiome was also prevalent in the water column and tank biofilm (Figure 2f-g). On closer observation, tank biofilm, tank water, and fish skin samples from the RAS systems were more similar compositionally than from the FT system (Figure 2, Supp Figure S3, Supp Figure S4), with RAS1 also having unique communities apart from RAS2.

**Figure 2.**
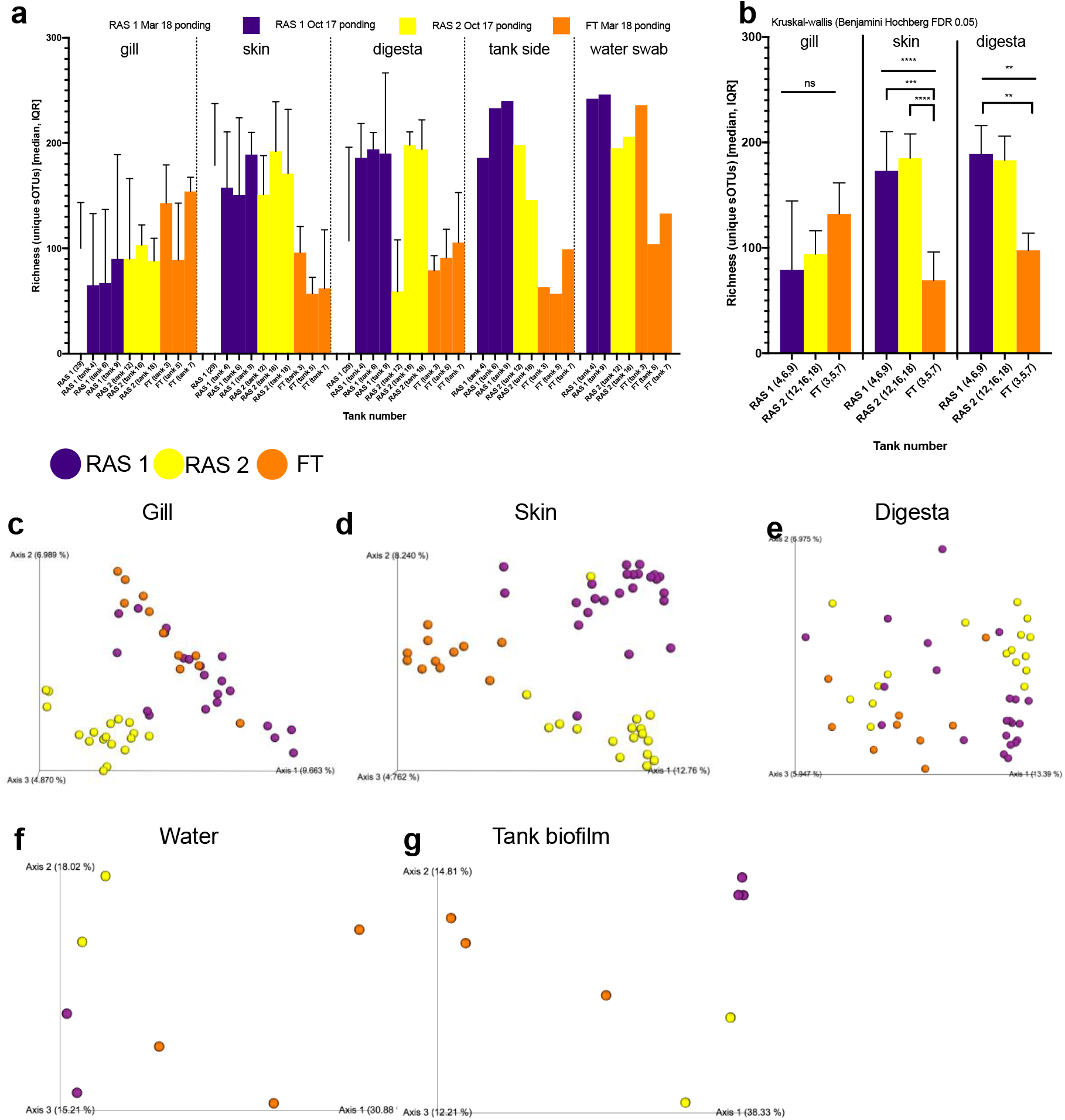
Inter-hatchery effects on microbial ecology of built environment and fish body sites. a) Richness (total observed sOTUs) distributions per each tank across body sites, tank biofilm, and water column from the three types of hatcheries RAS 1, RAS 2, and FT. b) Tank replicates are combined per hatchery to enable multiple group statistical analysis of richness comparisons (Kruskal-Wallis). Beta diversity distributions depicted through PCoA plots of Unweighted UniFrac distances across tanks for each unique environment: c) salmon gill, d) salmon skin, e) salmon digesta, f) tank water, and g) tank biofilm.

Next we directly evaluated the relationship between environmental microbiome of the tank water and tank biofilm with the fish mucus. For each individual tank, microbial richness of the biofilm (Figure 3a) and the tank water (Figure 3b) was compared to the richness of fish within that tank for the three body sites: gill, skin, and digesta. Both skin and digesta was positively correlated with tank biofilm (P=0.0001, R2 =0.2835; P=0.002, R2 =0.2042) and water richness (P=0.0014, R2 =0.2336; P=0.0264, R2 =0.1296) indicating that tank biofilms have a slightly stronger impact than tank water on fish mucus richness, with skin being the most impacted (Figure 3a-b). Since hatchery environmental microbes seemed to influence fish mucus microbes and unique microbial populations exist across hatcheries and within tank replicates within a hatchery, we hypothesized that within a tank, fish mucus microbial composition should be more similar to the biofilm and water of that tank as compared to tanks from other hatcheries. Here we report that the gill, skin, and digesta of the fish is more similar to the tank biofilm of origin compared to tanks from other hatcheries (Figure 3c) whereas for tank water this only is true for gill and skin (Figure 3d). Both gill and skin are more similar to the tank biofilm and tank water than digesta (Figure 3e). To understand how microbial communities differ across hatchery types, we compared the beta diversity within sample types from the three hatcheries. Tank water, tank biofilm, and skin communities are more similar between RAS hatcheries as compared to the FT hatchery (Supp Figure S4a-b). Since the skin microbiome was the most influenced body site on the fish, we calculated the differentially abundant sOTUs (n=65) between the RAS and FT systems (Supp Figure S4c). Of the 65 differentially abundant skin sOTUs, 44 were present in the water or tank biofilm communities, while 17 were only found on the skin (Supp Figure S4d). Skin microbes that were associated with RAS systems included *Saprospirales*, *Cytophagales*, *Sphingobacteriales*, *Verrucromicrobia*, and *Methylophilales* (*Methylotenera sp*), whereas the FT was enriched in *Pseudomonas*, *Pseudomonadales*, and *Enterobacteriales* (Supp Figure 4e). Additionally, Aeromonadales were highly enriched in the fecal detritus in the FT hatchery while much of the FT associated microbes were not found in the detritus suggesting they are indeed water or biofilm specific.

**Figure 3.**
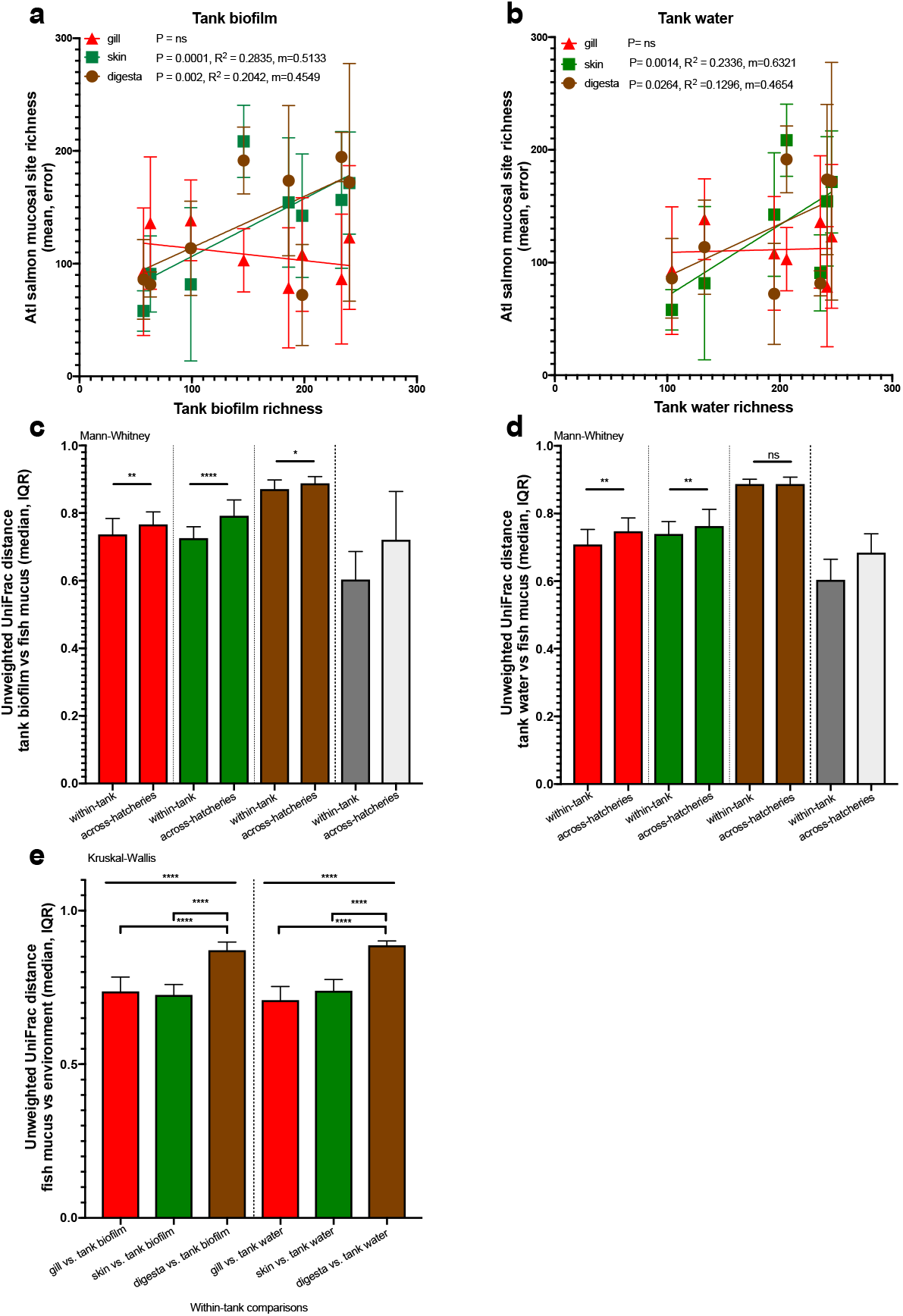
Relationship between built environment and fish mucosal microbiome. Correlation between a) tank biofilm richness and fish mucus richness along with b) tank water richness and fish mucus (linear regression). Beta diversity measures to test similarity (Unweighted UniFrac) of fish mucus to c) tank biofilm and d) tank water. Pairwise comparisons of similarities within a tank versus similarities to other tanks from across hatcheries with Mann-Whitney test. e) Overall fish mucosal similarities compared to tank biofilm and water indicate gill and skin are more similar to environment than digesta (Kruskal-Wallis)

**Figure 4.**
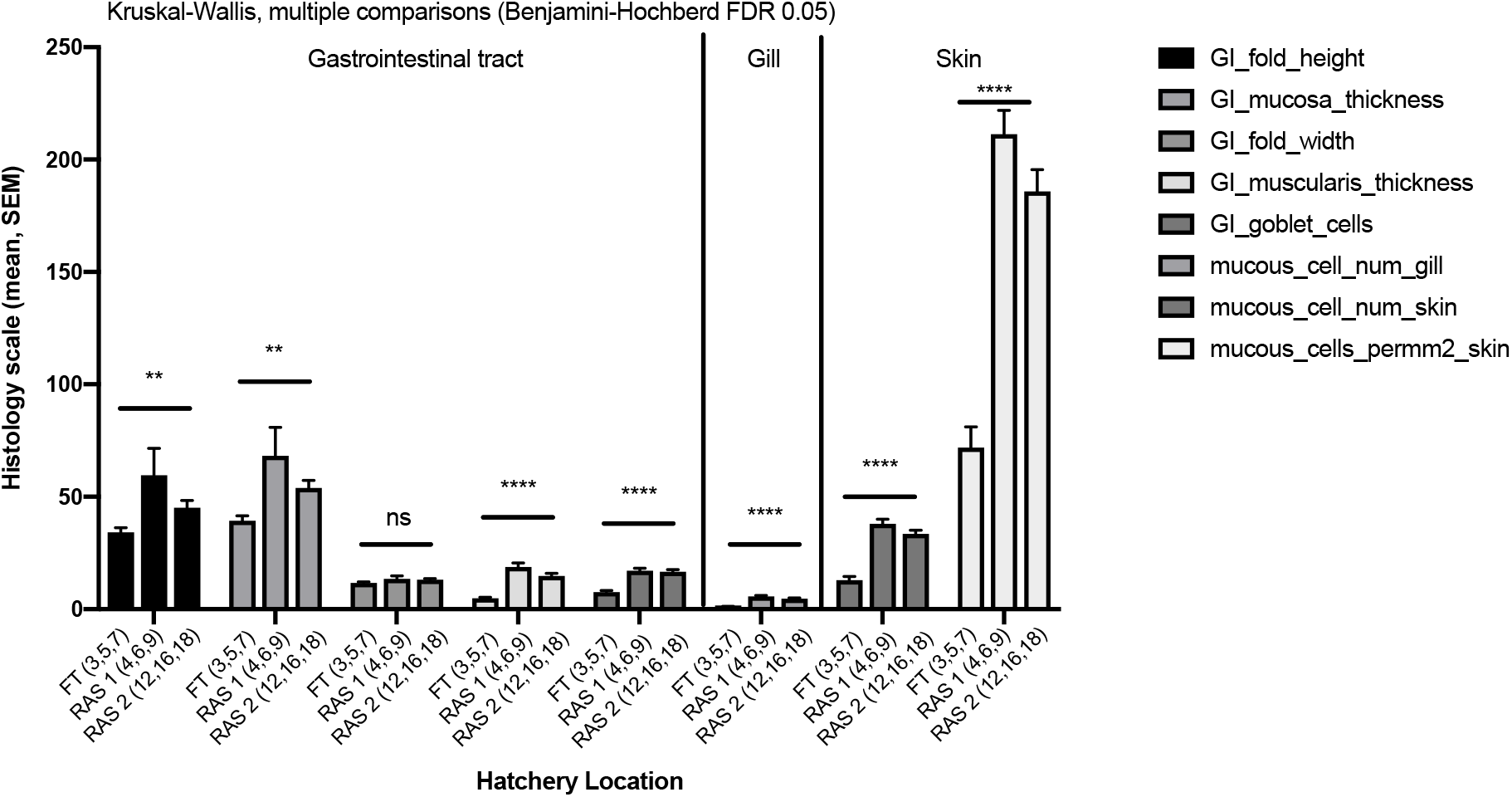
Histopathology analysis including gut morphometry and mucous cell counts from skin and gill of the fish from flow through (FT) and two RAS hatcheries (RAS 1 and RAS 2 hatchery systems. Hatchery systems were compared using non-parametric Kruskal-Wallis test. Skin mucous cell counts shown as both per length of epidermis section and per surface area or the epidermis. All fish sampled were clinically normal.

Upon establishing a direct relationship between the microbiome of the hatchery environment, we next assessed how fish health is related to these changes. Broad mucosal histopathology was performed on eight endpoint measures across the gill (Supp Figure S5), skin (Supp Figure S6), and gastrointestinal tract (Supp Figure S7). In all but one measure, a heighted score was demonstrated in RAS systems compared to FT for the fish sampled with RAS1 being slightly higher than RAS1 (Figure 4). Furthermore, we tested if the microbiome of the fish was driven by these histology scores and found that for Unweighted UniFrac measures, where rare taxa are more heavily weighted in a phylogenetic context, the skin microbiome was significantly associated with mucous cell numbers in the gill (Adonis: P=0.025) and skin communities (Adonis: P=0.006 and P=0003 while the gut microbiome was also associated with mucous cell numbers in the skin (Adonis: P=0.015) (Table 2). When analyzing weighted UniFrac, which looks primarily at relative abundances of sOTUs in a phylogenetic context, gill and skin microbial communities were associated with GI mucous cell numbers (Adonis: gill P=0.026, skin P=0.014) while the gut microbiome was associated with mucous cell numbers in the skin (Adonis: P=0.002) (Table 2).

**Table 2.**
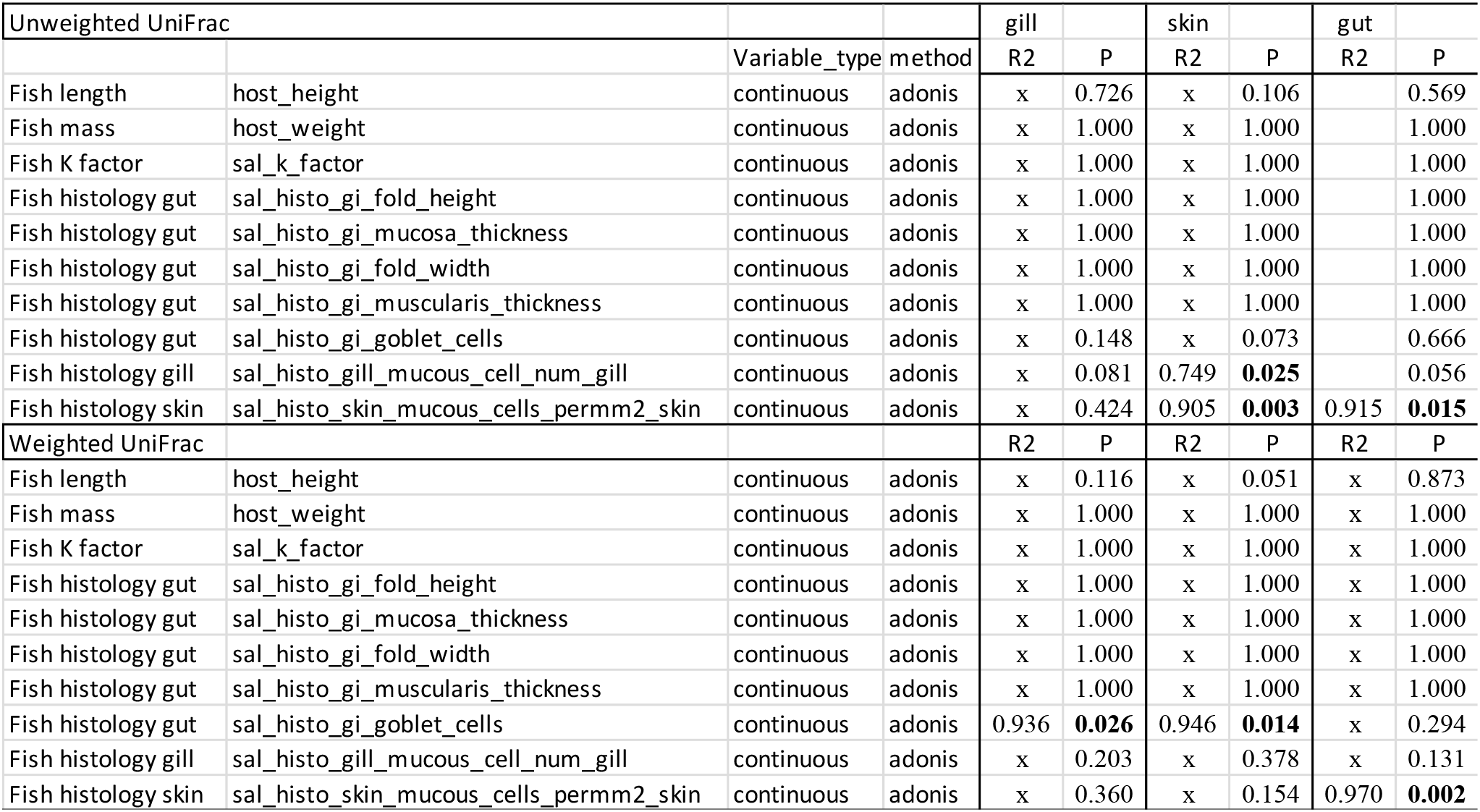
Multivariate statistical testing of effects of microbiome on fish health metrics

Recirculating Aquaculture Systems utilize microbes to recycle and remove nitrogenous waste products generated from uneaten feed, fish feces, and other organic wastes. We identified and quantified the types and relative abundances of these various types of known microbes (bacteria and archaea) in this system to understand if known RAS-associated microbes were playing a role in colonization within fish mucus or the environment (Figure 5a). The only known RAS-associated ammonia-oxidizing bacteria (AOB) found in the system was the family Nitrosomonadaceae which was present in all of the hatcheries and sample types (Figure 5b-e) and perhaps slightly enriched in the tank biofilm community (Figure 5f). Nitrite-oxidizing bacteria (NOB), primarily the family Nitrospiraceae and Nitrospira spp., were generally in higher relative abundances in the RAS environmental components including the water and biofilm (Figure 5e-f) along with the skin, digesta, and gill (Nitrospiraceae only) indicating a possible transfer event (Figure 5b-d). Note for digesta samples, both NOB organisms were not detected in any of the FT reared fish. For denitrifying autotrophs, Rhodobacter spp. and Hydrogenophaga spp. were enriched across all hatcheries and sample types with Rhodobacter spp. being in slightly higher abundances in some FT systems. For heterotrophic denitrifiers, Pseudomonas spp. were the most dominant and specifically were approximately 20-100× higher in the FT water and tank biofilms as compared to the RAS systems (Figure 5e-f). In addition to being enriched in the environment, Pseudomonas spp. were also consistently higher in the gill, skin, and digesta of fish reared in FT compared to RAS (Figure 5b-d). Lastly two methanogens were detected albeit at very low frequencies and only in the tank biofilm (Methanocorpusculum sp.) and the digesta (Methanosphaera sp.) from the RAS.

**Figure 5.**
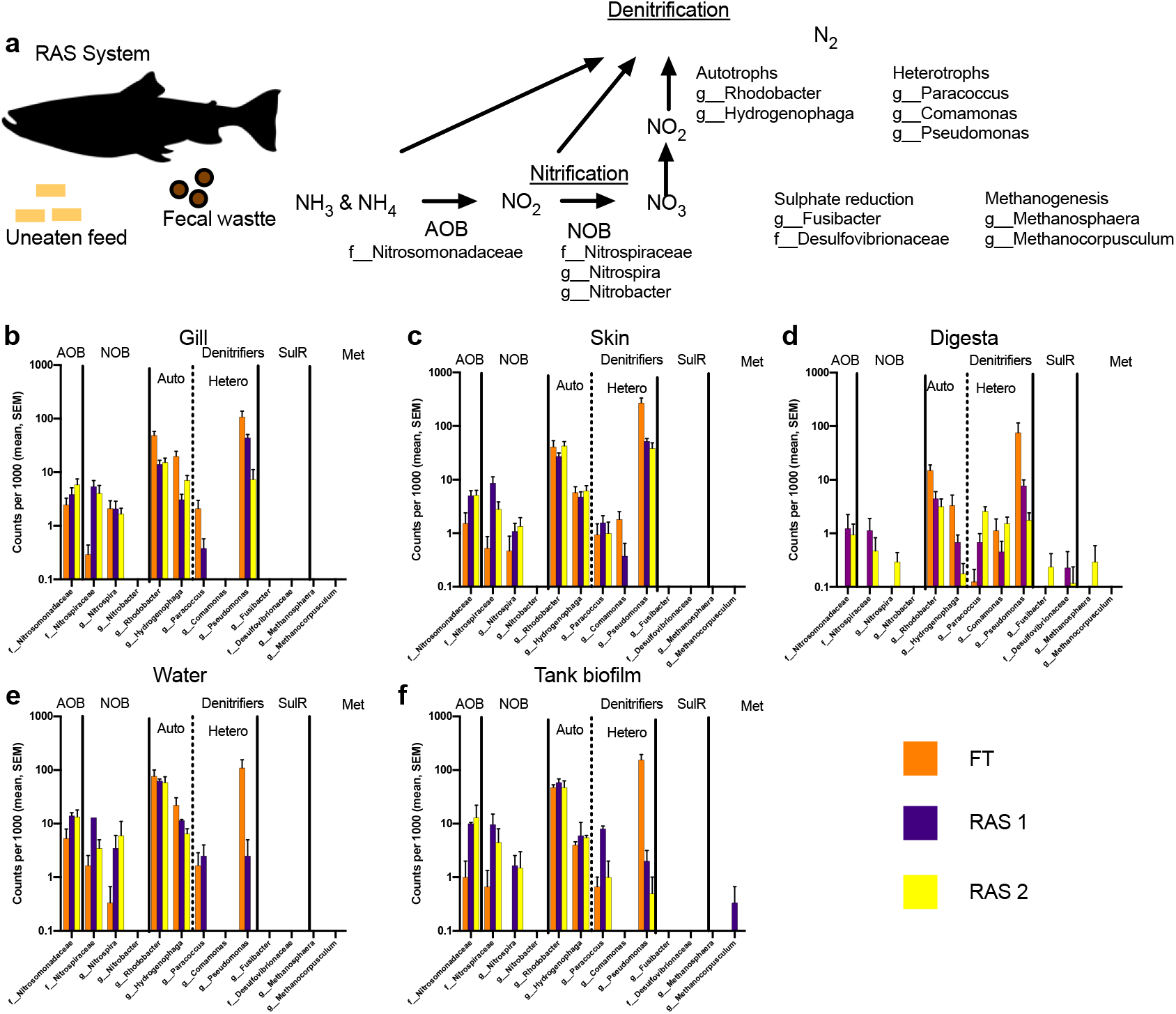
Distribution of RAS associated microbes in Salmon hatcheries. a) RAS systems are designed to recycle nitrogenous waste, primarily from uneaten feed and fish feces, using a series of nitrification and denitrification steps through microbial filters. The primary microbes involved in these processes and detected in the systems (AOB - ammonia oxidizing bacteria, NOB - Nitrite oxidizing bacteria, and denitrification) are listed. The distribution of these RAS-associated microbes are listed as mean relative counts per 1000 according to each hatchery type (FT = orange, RAS 1 = purple, RAS 2 = yellow) across each particular sample type including (b) gill, (c) skin, (d) digesta, (e) tank water, and (f) tank biofilm.

## DISCUSSION

The mucosal environment is paramount for fish health as it is the first line of defense against pathogen invasion. Specifically, a healthy mucosal environment protects against infection through several endogenous mechanisms including mucus production, immune components such as lysozymes, antimicrobial peptides, immunoglobulins, and exogenous mechanisms through establishment of a healthy microbiome. In this study, we investigated the means by which the mucosal environment of Atlantic salmon is influenced by the rearing environment. We evaluated three unique hatcheries utilizing two rearing methodologies including Recirculating Aquaculture Systems (RAS) and Flow-through (FT) systems.

In both the biosecure RAS and FT hatchery environments, Atl salmon have unique microbial communities on their gill, skin, and digesta. These fish associated mucosal microbiomes along with the tank and biofilm communities are further differentiated across hatchery systems by comparing RAS vs. FT systems. RAS systems are known to harbor their own unique microbial communities both in the biofilter but also within the hatchery system where fish are reared (47), Previous studies however, have not looked at the built environment microbiomes simultaneously with the fish mucosal microbiomes. For these hatchery systems, alpha diversity is higher in RAS compared to FT hatcheries for the following sample types: skin, digesta, tank water, and tank biofilm microbiomes. Fish skin and digesta richness is further positively associated with both tank biofilm and tank water richness suggesting an influence of the environment microbiota on fish associated microbiota, with the biofilm association being the strongest. Skin microbiomes have been implicated as important for maintaining fish health, thus understanding any potential negative implications or drivers of dysbiosis is important for fish welfare (48, 49). Tank biofilms can be challenging to monitor and control. Further research should focus on how manipulating tank surfaces through material science and engineering could be used to promote fish health.

Beta-diversity is significantly different across the three hatcheries when looking at individual sample types: gill, skin, digesta, water, and biofilm. Fish mucosal sites were more phylogenetically similar to both water and biofilms within their own tank as compared to tanks from other hatcheries indicating a microenvironment effect. By performing histology of fish GI, skin, and gill we confirmed that the fish mucosal microbiome is associated with fish health.

RAS are becoming popular for growing salmon smolts offering many benefits including minimized water use and waste generation along with improving survival rates of fish during transfer to net pens (50–53). Waste water is purified by processing through one of two main types of biofilters (fixed film or single sludge) which utilizes a variety of bacteria and archaea (54, 55). The biofilters are primarily comprised of heterotrophs and chemoautotrophs that transform and detoxify ammonia and nitrate species (56). The common ammonia-oxidizing archaea and bacteria found in these systems include *Nitrosopumilus* (archaea), *Nitrosomonas*, *Nitrosococcus*, and *Nitrosospira* (47). In our study, only ammonia-oxidizing bacteria within the family Nitrosomonadaceae were present and were highest in the RAS tank and water systems as well as RAS reared fish gill, skin, and digesta. Following ammonia oxidation, *Nitrospira* and *Nitrobacter* are the primary bacteria responsible for nitrite oxidation in RAS biofilters (47). Bacterial sOTUs from *Nitrospira* sp and unclassified sOTUs within the family Nitrospiraceae were in higher relative abundance in the RAS hatcheries for tank water, tank biofilm, skin, digesta, and moderately in the gill. In the final step of nitrogen recycling, denitrification is carried out by both autotrophs and heterotrophs. The primary autotrophic bacteria associated with denitrification in RAS systems include *Thiomicrospira*, *Thiothrix*, *Rhodobacter*, and *Hydrogenophaga* (47). Both *Rhodobacter* and *Hydrogenophaga* were found in the hatcheries although in similar relative abundances across the FT and RAS hatcheries both in the tank environment and fish mucus. The primary Heterotrphic microbes associated with denitrification in RAS systems includes *Pseudomonas*, *Paracoccus*, and *Comamonas sp* (47). All three were abundant in the hatcheries with Pseudomonas being the highest of the three and generally higher in the flow through hatchery compared to RAS. In conclusion, various RAS associated microbes which are responsible for Nitrogen cycling, particularly Nitrification, in the biofilters were present in our study and higher in the RAS built environment along with RAS reared fish mucus suggesting that these microbes are not being solely sequestered in the biofilter but instead also are circulated through the fish tanks and may be colonizing fish mucus.

When excess organic matter including fish feed and fish feces accumulates in a RAS tank, the heterotrophs can quickly bloom and outcompete nitrifying microbes (47). This overgrowth and imbalance may contribute negatively to flesh flavor, thus future studies are warranted to understand which microbes and what metabolic pathways may play this role (57). While most hatcheries are used for producing seed to then transfer to ocean growout cages, complete salmon production cycles in land based RAS is becoming more common. Furthermore, both FT and RAS systems may be colonized by various microbial inputs from the air, water, fish feed, fish flesh, technicians, and biofilter type, thus understanding the contributions of each in a system will be important both for future experimental designs and for fish health (58).

The built environment microbiome may originally be colonized by both animal excrement including mucus along with environmental sources such as water. The sustained built environment microbiome is both a result of the new animal host deposition of cellular material but can also propagate based on host associated animal matter. Furthermore, the built environment community can then influence the microbial communities of animal hosts residing there. Understanding the extent by which the animal’s microbiome can be influenced by its surroundings and then associated to a phenotype such as fish health or development will be important for experimental design where microbiome readout is a standard measure. This is commonly referred to as the ‘cage effect’ and has primarily been demonstrated in mouse studies where animals which share the same cage have more similar fecal samples, likely due to coprophagy (59, 60). Cage effect can explain up to 31% of variation in mouse feces compared to only 19% resolved by host genetics (61). For this reason, our experimental design included three separate tanks per treatment group (hatchery) along with multiple fish biological replicates per tank. To our knowledge, this is the first experiment to demonstrate a tank effect in fish which is due to both water and tank biofilm formation and influences primarily the fish skin and digesta. Since aquariums use a variety of material types to culture fish, it would be important for future studies to evaluate how biofilm formation changes with respect to tank material type (e.g. concrete, PVC, HDPE, fiberglass, etc).

Quantifying fish health can be a challenging and expensive endeavor which does not often easily scale for large hatchery operations. Fish mucus contains various immune components such as lysozymes, immunoglobins, lectins, crinotoxins, and antimicrobial peptides (62). Here we used histology as a measure of mucosal health across the gastrointestinal tract, gill, and skin. Elevated skin mucous cell numbers are generally reflective of healthy fish whereas depleted mucosal cells may indicate a recent mucosal discharge due to stress or disease (63, 64). Mucosal cell numbers however, can also be influenced by the sampling location on the fish body, sex, diet, and age or development stage, thus care must be taken when interpreting results (31, 65, 66). In our study, skin and gill mucosal cell numbers were elevated in both RAS systems compared to the FT suggesting that RAS fish may have been more healthy or less stressed. Furthermore, these elevated skin mucous cell numbers positively correlated to microbial richness and phylogenetic diversity on the skin and were associated with changes in microbial composition. For the gastrointestinal tract, the fold height, mucosa thickness, muscularis thickness, and mucous cell number were higher in fish reared in the RAS compared to the FT hatchery. Overall, the RAS reared fish had a more complex GI tract compared to fish reared in flow through systems. Vertebrate gut microbiomes are often driven by diet, habitat, and age (67). Since diet and age was controlled for in this study, we hypothesize that differences in microbial communities in the tank water column and biofilm are driving gut microbiome differences by drinking or grazing. Microbiome is essential for development and differentiation of mucous cells, for example gnobiotic model of zebrafish showed reduced numbers of mucous cells in their intestine (Bates et al 2006). For each unique mucosal site, distinct microbial communities were present and differentiated between the FT and RAS systems. Differences in GI communities between RAS and open water systems may be indicative of microbial exposure in the environment (68). Atlantic salmon reared in a RAS which were infected by *Aeromonas salmonicida,* also had differentiated gut microbiomes as compared to healthy fish (69). By demonstrating how the environmental microbiome is influenced by hatchery design which in turn influences the fish mucosal microbiome and subsequent health, our study demonstrates the utility of developing environmental and/or fish microbiome sampling as a potential fish tank health indicator. Future studies should evaluate more tanks and include metrics such as survival rate, growth rate, and body composition analysis to determine how the environmental microbiome may drive fish performance in the hatchery setting.

---

**Supplemental Figure S1.**
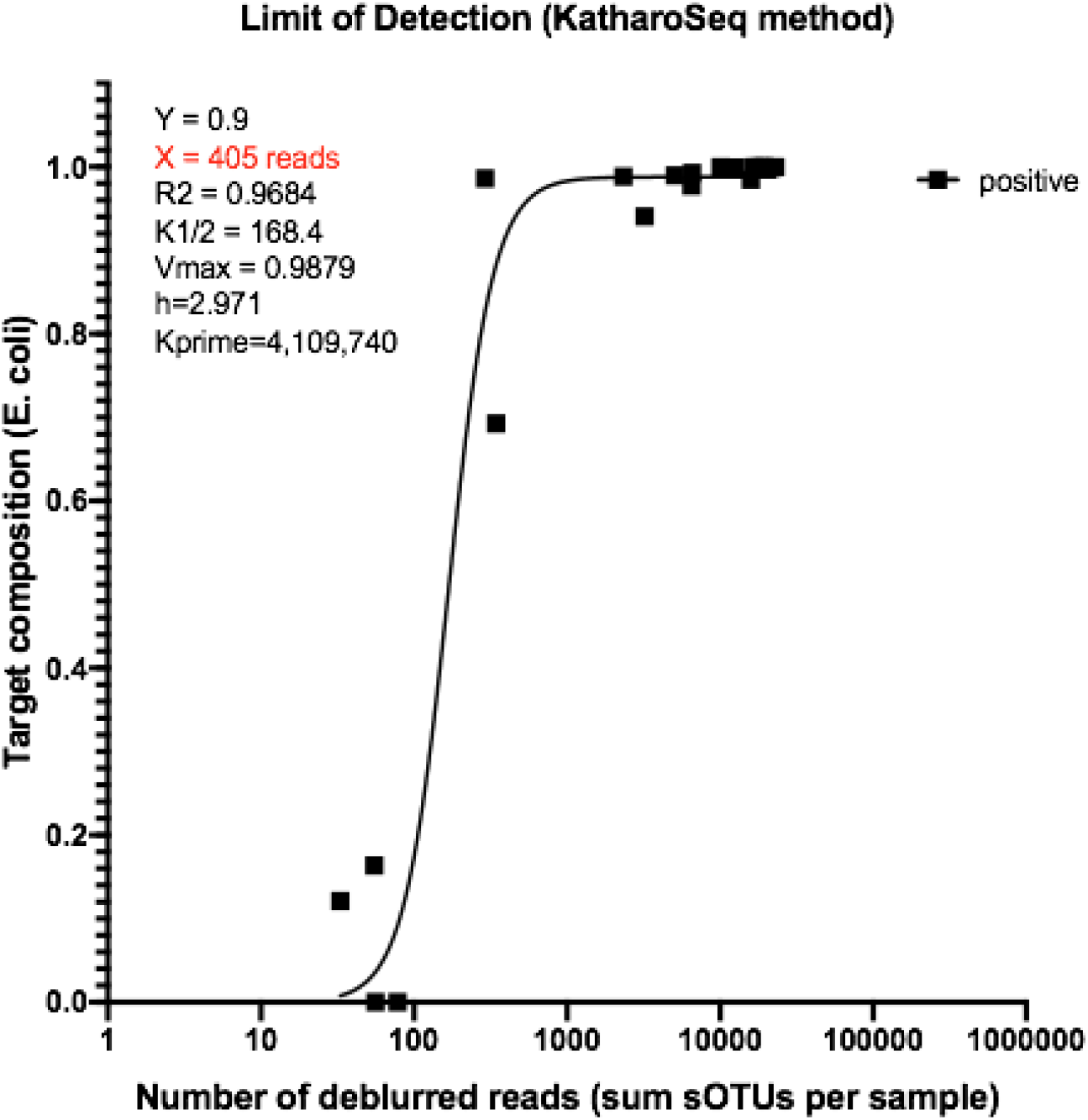
Limit of detection calculation of positive control titrations. Application of Katharoseq method results in a cutoff value of 405 reads indicating that positive control samples which have 405 reads would then have 90% of those reads aligning to the target organism.

**Supplemental Figure S2.**
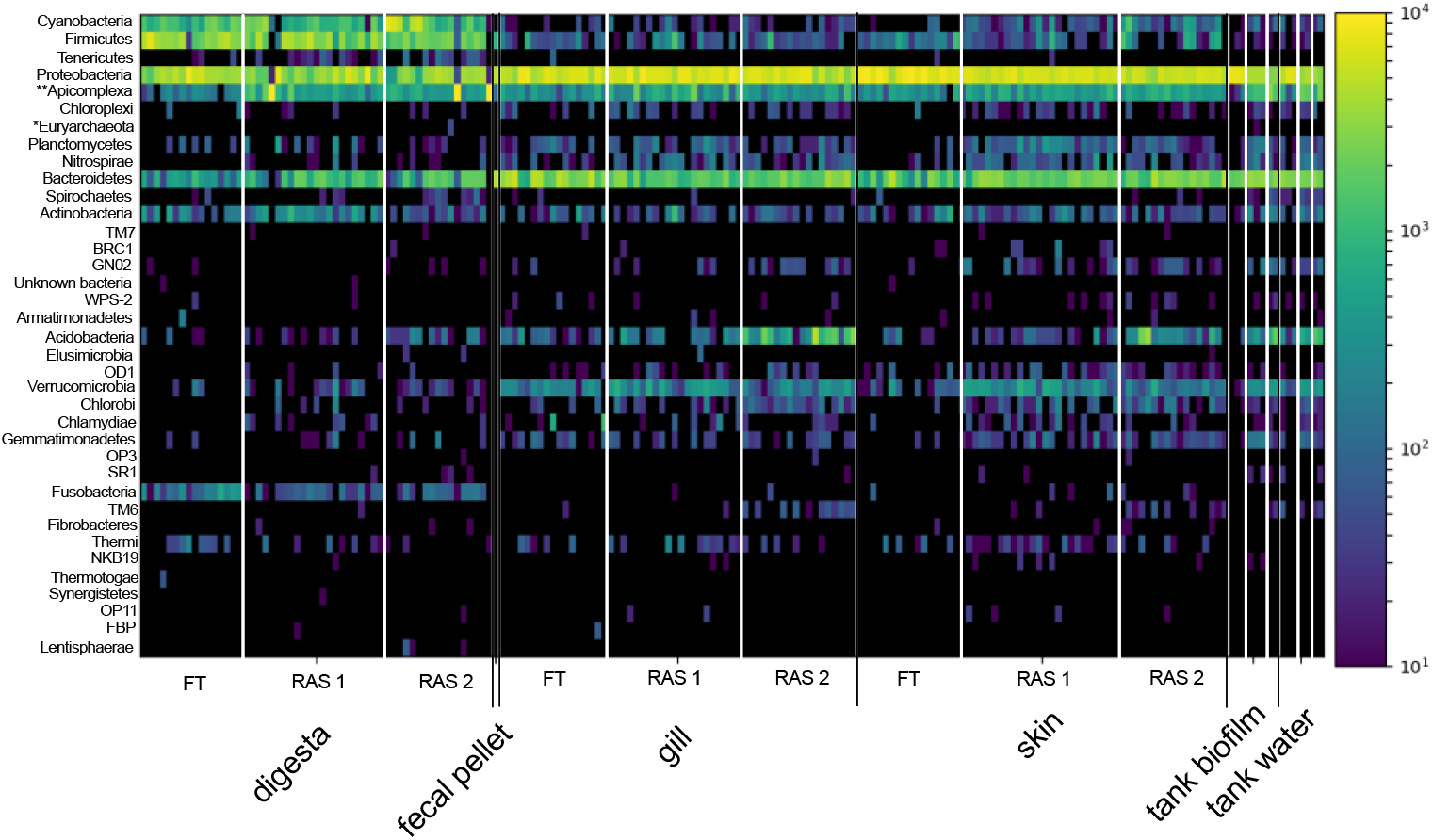
Microbial phyla (37) present in the study including bacteria, *archea, and ** one microbial eukaryote.

**Supplemental Figure S3.**
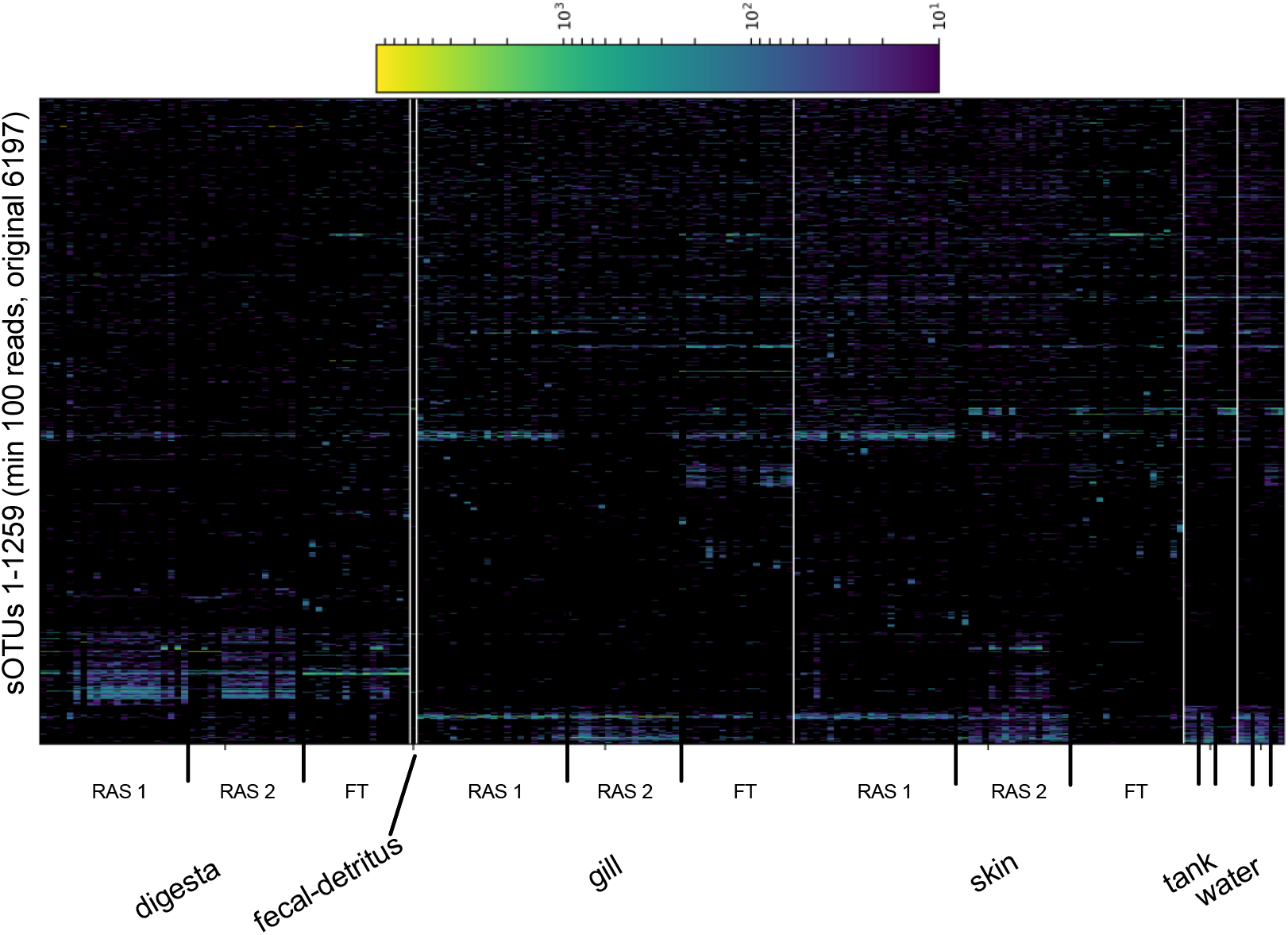
Top 20% most abundant sOTUs (minimum 100 reads across sample types)

**Supplemental Figure S4.**
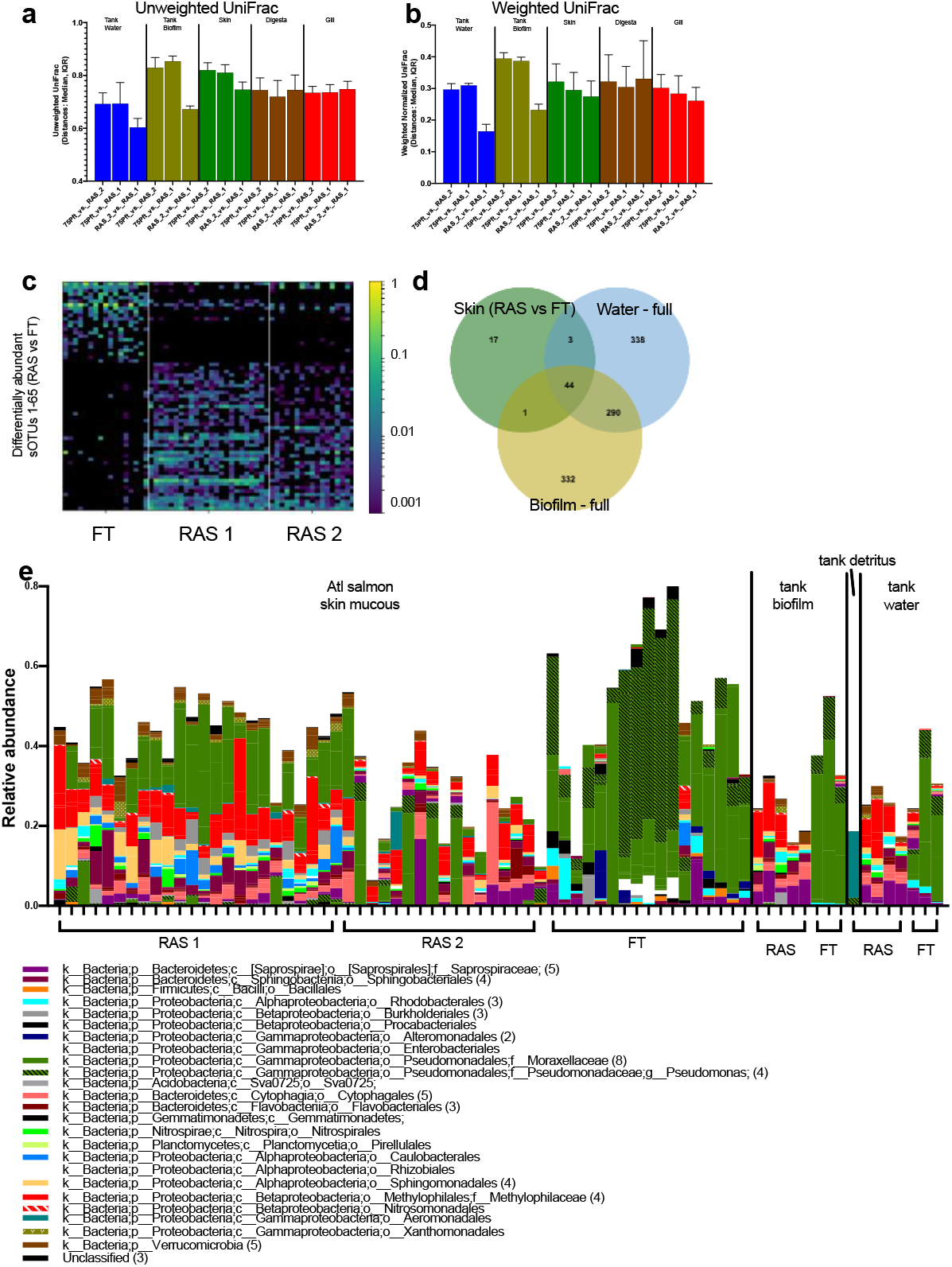
Microbial differences between hatchery type. Beta-diversity comparisons between all three hatcheries (FT, RAS 1, and RAS 2) for individual sample types including tank water, tank biofilm, skin, digesta, and gill for a) Unweighted Unifrac and b) Weighted UniFrac. c) Comparing skin microbiomes of fish from the RAS vs. the FT hatcheries, 65 sOTUs were differentially abundant. d) Source tracking of differentially abundant skin microbes from water and tank biofilm communities demonstrate that the majority are founds in built environment. e) Differentially abundant skin microbes colored by taxonomic order found across the three hatcheries and abundances within the tank water and tank biofilm.

**Supplemental Figure S5.**
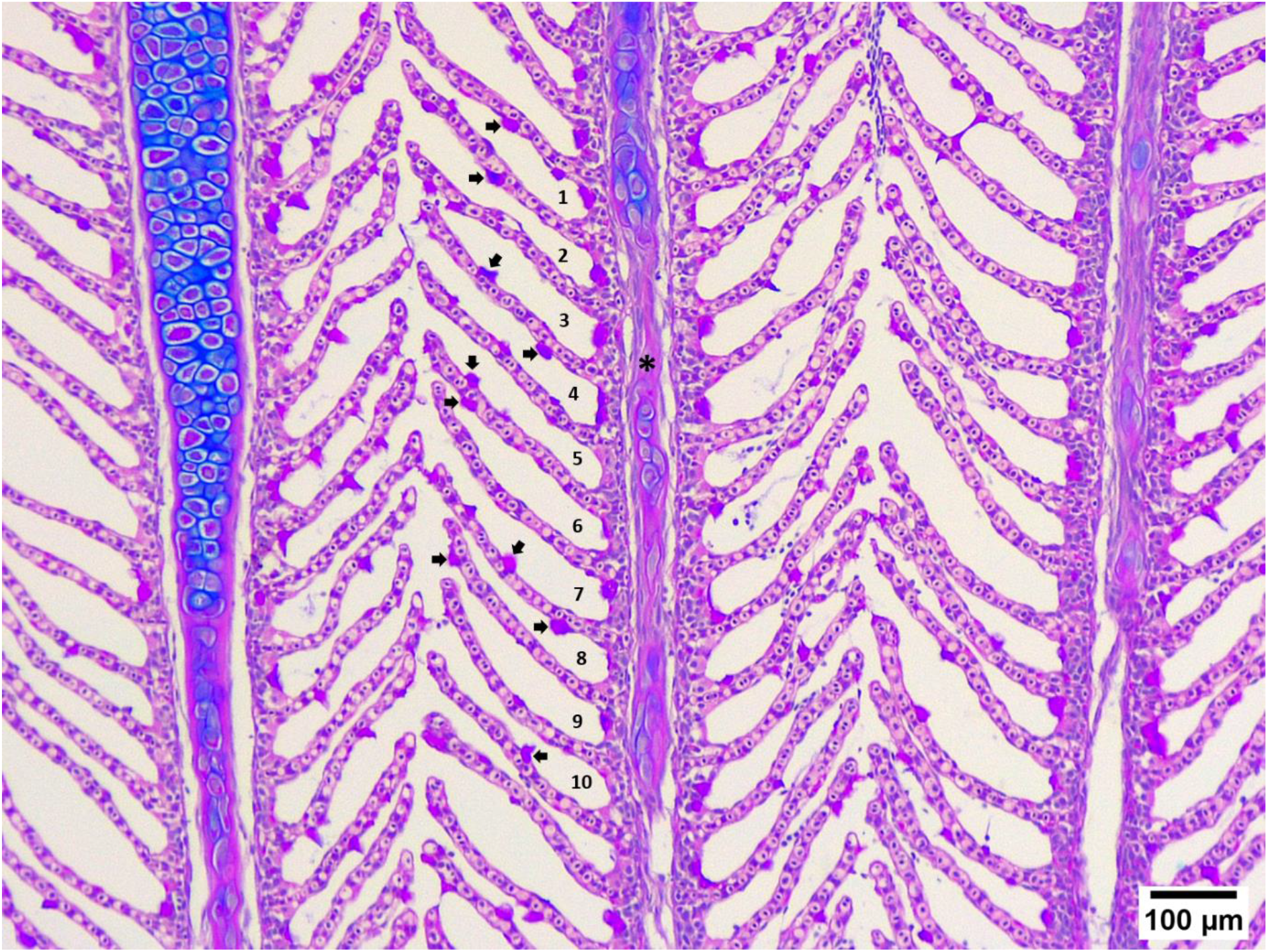
Atlantic salmon’s gills stained with Alcian Blue/ Periodic Acid – Schiff (AB/PAS) at pH 2.5 illustrating the mucous cells (arrow) in gill lamellae of a well-orientated gill filament (*). The gill mucous cells were quantified in ten gills interlamellar unit (ILU) for each sample under the light microscope. Scale bar 100 μm.

**Supplemental Figure S6.**
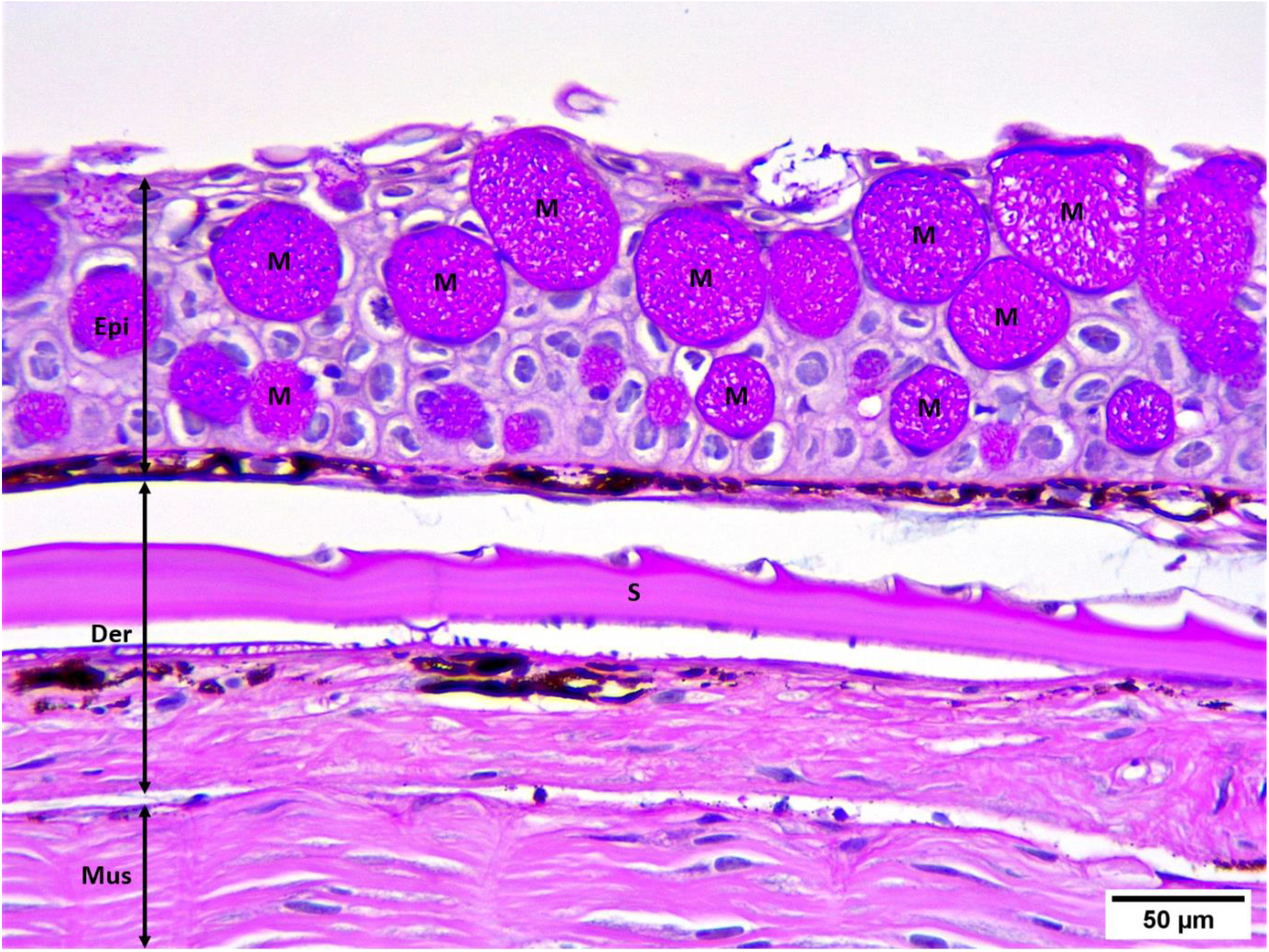
Atlantic salmon’s skin section stained with Alcian Blue/ Periodic Acid–Schiff (AB/PAS) at pH 2.5 showing the mucous cells (M) containing mixed mucin, purple in colour, in the epidermis (Epi). The skin mucous cells were counted in 5 random areas of each fish sample with the same magnification (S=scale, Der=dermis, Mus=muscle). Scale bar 50 μm.

**Supplemental Figure S7.**
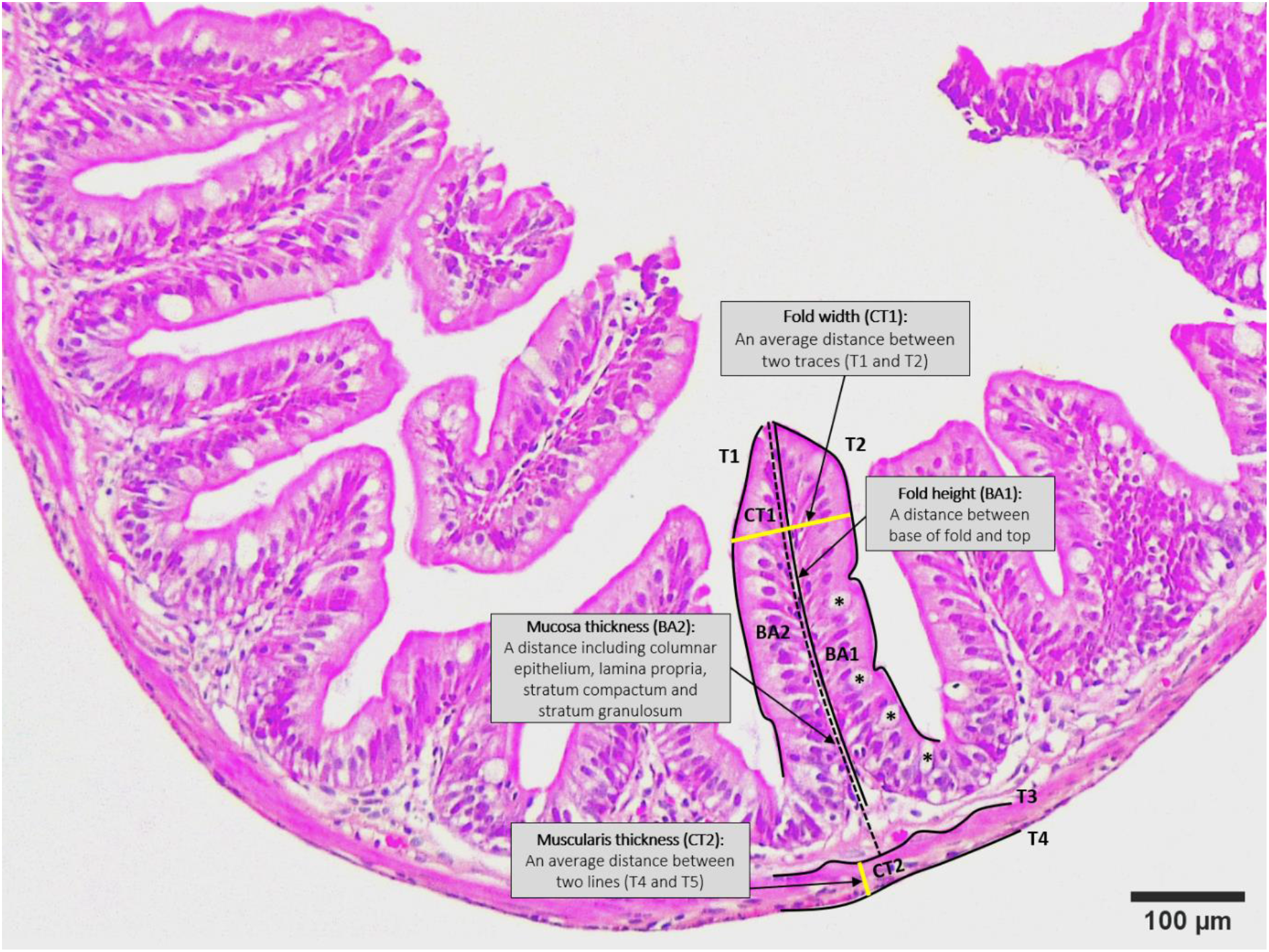
Atlantic salmon’s intestine section stained with haematoxylin and eosin (H&E) demonstrating the general organization of the intestinal wall. Vertical and horizontal lines show how the fold height (BA1), mucosa thickness (BA2), fold width (CT1), and muscularis thickness (CT2) were measured in the morphometrical analyses using Image-Pro Premier 9.1 software. The intestinal mucous cells (*) in the measured fold were counted under the light microscope while capturing images for the gut morphometrical analyses. Scale bar 100 μm.

## REFERENCES

1. FAO. 2016. The State of World Fisheries and Aquaculture 2016 (SOFIA): Contributing to food security and nutrition for all. FAO, Rome, Italy.

2. Gentry RR, Froehlich HE, Grimm D, Kareiva P, Parke M, Rust M, Gaines SD, Halpern BS. 2017. Mapping the global potential for marine aquaculture. Nat Ecol Evol 1:1317–1324.

3. 2018. Meeting the sustainable development goals. Rome.

4. Tave D, Hutson AM. Is Good Fish Culture Management Harming Recovery Efforts in Aquaculture-Assisted Fisheries? North Am J Aquac 0.

5. Kumar G, Engle CR. 2016. Technological Advances that Led to Growth of Shrimp, Salmon, and Tilapia Farming. Rev Fish Sci Aquac 24:136–152.

6. Stopha M. Alaska salmon fisheries enhancement annual report 2018. 92.

7. Jonsson B, Jonsson N. 2006. Cultured Atlantic salmon in nature: a review of their ecology and interaction with wild fish. ICES J Mar Sci 63:1162–1181.

8. Jonsson N, Jonsson B, Hansen LP. 2003. The marine survival and growth of wild and hatchery-reared Atlantic salmon. J Appl Ecol 40:900–911.

9. Jensen AJ, Berg M, Bremset G, Finstad B, Hvidsten NA, Jensås JG, Johnsen BO, Lund E. 2016. Passing a seawater challenge test is not indicative of hatchery-reared Atlantic salmon Salmo salar smolts performing as well at sea as their naturally produced conspecifics. J Fish Biol 88:2219–2235.

10. d’Orbcastel ER, Blancheton J-P, Aubin J. 2009. Towards environmentally sustainable aquaculture: Comparison between two trout farming systems using Life Cycle Assessment. Aquac Eng 40:113–119.

11. Eding EH, Kamstra A, Verreth JAJ, Huisman EA, Klapwijk A. 2006. Design and operation of nitrifying trickling filters in recirculating aquaculture: A review. Aquac Eng 34:234–260.

12. Badiola M, Basurko OC, Piedrahita R, Hundley P, Mendiola D. 2018. Energy use in Recirculating Aquaculture Systems (RAS): A review. Aquac Eng 81:81–57.

13. Attramadal KJK, Truong TMH, Bakke I, Skjermo J, Olsen Y, Vadstein O. 2014. RAS and microbial maturation as tools for K-selection of microbial communities improve survival in cod larvae. Aquaculture 432:432–483.

14. Martins CIM, Pistrin MG, Ende SSW, Eding EH, Verreth JAJ. 2009. The accumulation of substances in Recirculating Aquaculture Systems (RAS) affects embryonic and larval development in common carp Cyprinus carpio. Aquaculture 291:291–65.

15. Deviller G, Palluel O, Aliaume C, Asanthi H, Sanchez W, Franco Nava MA, Blancheton J-P, Casellas C. 2005. Impact assessment of various rearing systems on fish health using multibiomarker response and metal accumulation. Ecotoxicol Environ Saf 61:61–89.

16. Martins C, Ende S, Ochola D, Eding E, Verreth J. 2007. Growth retardation in Nile tilapia (Oreochromis niloticus) cultured in recirculation aquaculture systems, p. 162–163. *In*.

17. Bakke I, Åm AL, Kolarevic J, Ytrestøyl T, Vadstein O, Attramadal KJK, Terjesen BF. 2017. Microbial community dynamics in semi-commercial RAS for production of Atlantic salmon post-smolts at different salinities. Aquac Eng 78:78–42.

18. Beck BH, Peatman E. 2015. Mucosal Health in Aquaculture. Academic Press.

19. Adams MB, Nowak BF. 2003. Amoebic gill disease: sequential pathology in cultured Atlantic salmon, Salmo salar L. J Fish Dis 26:26–601.

20. Llewellyn MS, Leadbeater S, Garcia C, Sylvain F-E, Custodio M, Ang KP, Powell F, Carvalho GR, Creer S, Elliot J, Derome N. 2017. Parasitism perturbs the mucosal microbiome of Atlantic Salmon. Sci Rep 7:43465.

21. Bakke-McKellep AM, Press CM, Baeverfjord G, Krogdahl Å, Landsverk T. 2000. Changes in immune and enzyme histochemical phenotypes of cells in the intestinal mucosa of Atlantic salmon, Salmo salar L., with soybean meal-induced enteritis. J Fish Dis 23:23–115.

22. Webster TMU, Consuegra S, Hitchings M, Leaniz CG de. 2018. Interpopulation Variation in the Atlantic Salmon Microbiome Reflects Environmental and Genetic Diversity. Appl Environ Microbiol 84:e00691–18.

23. Lokesh J, Kiron V. 2016. Transition from freshwater to seawater reshapes the skin-associated microbiota of Atlantic salmon. Sci Rep 6:19707.

24. Llewellyn MS, McGinnity P, Dionne M, Letourneau J, Thonier F, Carvalho GR, Creer S, Derome N. 2016. The biogeography of the atlantic salmon (*Salmo salar*) gut microbiome. ISME J 10:10–1280.

25. Hovda MB, Fontanillas R, McGurk C, Obach A, Rosnes JT. 2012. Seasonal variations in the intestinal microbiota of farmed Atlantic salmon (Salmo salar L.): Seasonal variations in the intestinal microbiota of Salmo salar L. Aquac Res 43:43–154.

26. Dehler CE, Secombes CJ, Martin SAM. 2017. Seawater transfer alters the intestinal microbiota profiles of Atlantic salmon (Salmo salar L.). Sci Rep 7:13877.

27. Minich JJ, Zhu Q, Janssen S, Hendrickson R, Amir A, Vetter R, Hyde J, Doty MM, Stillwell K, Benardini J, Kim JH, Allen EE, Venkateswaran K, Knight R. 2018. KatharoSeq Enables High-Throughput Microbiome Analysis from Low-Biomass Samples. mSystems 3:e00218–17.

28. Gilbert JA, Stephens B. 2018. Microbiology of the built environment. Nat Rev Microbiol 16:16–661.

29. Dang M, Nørregaard R, Bach L, Sonne C, Søndergaard J, Gustavson K, Aastrup P, Nowak B. 2017. Metal residues, histopathology and presence of parasites in the liver and gills of fourhorn sculpin (Myoxocephalus quadricornis) and shorthorn sculpin (Myoxocephalus scorpius) near a former lead-zinc mine in East Greenland. Environ Res 153:153–171.

30. Roberts SD, Powell MD. 2003. Comparative ionic flux and gill mucous cell histochemistry: effects of salinity and disease status in Atlantic salmon (Salmo salar L.). Comp Biochem Physiol A Mol Integr Physiol 134:134–525.

31. Pittman K, Sourd P, Ravnøy B, Espeland Ø, Fiksdal IU, Oen T, Pittman A, Redmond K, Sweetman J. 2011. Novel method for quantifying salmonid mucous cells: Quantification of salmonid mucous cells. J Fish Dis 34:34–931.

32. L⊘kka G, Austb⊘ L, Falk K, Bjerkås I, Koppang EO. 2013. Intestinal morphology of the wild atlantic salmon (Salmo salar). J Morphol 274:274–859.

33. Minich JJ, Sanders JG, Amir A, Humphrey G, Gilbert JA, Knight R. 2019. Quantifying and Understanding Well-to-Well Contamination in Microbiome Research. mSystems 4.

34. Minich JJ, Humphrey G, Benitez RAS, Sanders J, Swafford A, Allen EE, Knight R. 2018. High-Throughput Miniaturized 16S rRNA Amplicon Library Preparation Reduces Costs while Preserving Microbiome Integrity. mSystems 3:e00166–18.

35. Parada AE, Needham DM, Fuhrman JA. 2016. Every base matters: assessing small subunit rRNA primers for marine microbiomes with mock communities, time series and global field samples. Environ Microbiol 18:18–1403.

36. Walters W, Hyde ER, Berg-Lyons D, Ackermann G, Humphrey G, Parada A, Gilbert JA, Jansson JK, Caporaso JG, Fuhrman JA, Apprill A, Knight R. 2016. Improved Bacterial 16S rRNA Gene (V4 and V4-5) and Fungal Internal Transcribed Spacer Marker Gene Primers for Microbial Community Surveys. mSystems 1:e00009–15.

37. Caporaso JG, Lauber CL, Walters WA, Berg-Lyons D, Huntley J, Fierer N, Owens SM, Betley J, Fraser L, Bauer M, Gormley N, Gilbert JA, Smith G, Knight R. 2012. Ultra-high-throughput microbial community analysis on the Illumina HiSeq and MiSeq platforms. ISME J 6:1621–1624.

38. Gonzalez A, Navas-Molina JA, Kosciolek T, McDonald D, Vázquez-Baeza Y, Ackermann G, DeReus J, Janssen S, Swafford AD, Orchanian SB, Sanders JG, Shorenstein J, Holste H, Petrus S, Robbins-Pianka A, Brislawn CJ, Wang M, Rideout JR, Bolyen E, Dillon M, Caporaso JG, Dorrestein PC, Knight R. 2018. Qiita: rapid, web-enabled microbiome meta-analysis. Nat Methods 15:15–796.

39. Caporaso JG, Kuczynski J, Stombaugh J, Bittinger K, Bushman FD, Costello EK, Fierer N, Peña AG, Goodrich JK, Gordon JI, Huttley GA, Kelley ST, Knights D, Koenig JE, Ley RE, Lozupone CA, McDonald D, Muegge BD, Pirrung M, Reeder J, Sevinsky JR, Turnbaugh PJ, Walters WA, Widmann J, Yatsunenko T, Zaneveld J, Knight R. 2010. QIIME allows analysis of high-throughput community sequencing data. Nat Methods 7:7–335.

40. Amir A, McDonald D, Navas-Molina JA, Kopylova E, Morton JT, Zech Xu Z, Kightley EP, Thompson LR, Hyde ER, Gonzalez A, Knight R. 2017. Deblur Rapidly Resolves Single-Nucleotide Community Sequence Patterns. mSystems 2.

41. Kruskal WH, Wallis WA. 1952. Use of Ranks in One-Criterion Variance Analysis. J Am Stat Assoc 47:47–583.

42. Benjamini Y, Hochberg Y. 1995. Controlling the False Discovery Rate: A Practical and Powerful Approach to Multiple Testing. J R Stat Soc Ser B Methodol 57:57–289.

43. Lozupone C, Knight R. 2005. UniFrac: a New Phylogenetic Method for Comparing Microbial Communities. Appl Environ Microbiol 71:71–8228.

44. Chang Q, Luan Y, Sun F. 2011. Variance adjusted weighted UniFrac: a powerful beta diversity measure for comparing communities based on phylogeny. BMC Bioinformatics 12:118.

45. Anderson MJ. 2001. A new method for non-parametric multivariate analysis of variance. Austral Ecol 26:26–32.

46. Xu ZZ, Amir A, Sanders J, Zhu Q, Morton JT, Bletz MC, Tripathi A, Huang S, McDonald D, Jiang L, Knight R. 2019. Calour: an Interactive, Microbe-Centric Analysis Tool. mSystems 4:e00269–18.

47. Rurangwa E, Verdegem MCJ. 2015. Microorganisms in recirculating aquaculture systems and their management. Rev Aquac 7:7–117.

48. Legrand TPRA, Catalano SR, Wos-Oxley ML, Stephens F, Landos M, Bansemer MS, Stone DAJ, Qin JG, Oxley APA. 2018. The Inner Workings of the Outer Surface: Skin and Gill Microbiota as Indicators of Changing Gut Health in Yellowtail Kingfish. Front Microbiol 8.

49. Merrifield DL, Rodiles A. 2015. 10 - The fish microbiome and its interactions with mucosal tissues, p. 273–295. *In* Beck, BH, Peatman, E (eds.), Mucosal Health in Aquaculture. Academic Press, San Diego.

50. Terjesen B, Ulgenes Y, Fjæra S, Summerfelt S, Brunsvik P, Baeverfjord G, Nerland S, Takle H, Norvik O, Kittelsen A. 2008. RAS research facility dimensioning and design: A special case compared to planning production systems, p. 223–238. *In*. Citeseer.

51. Bergheim A, Drengstig A, Ulgenes Y, Fivelstad S. 2009. Production of Atlantic salmon smolts in Europe—current characteristics and future trends. Aquac Eng 41:41–46.

52. Drengstig A, Ulgenes Y, Liltved H, Bergheim A. 2011. Recent RAS trends of commercial salmon smolt farming in Norway. Aquac Am.

53. Martins C, Eding EH, Verdegem MC, Heinsbroek LT, Schneider O, Blancheton J-P, d’Orbcastel ER, Verreth J. 2010. New developments in recirculating aquaculture systems in Europe: A perspective on environmental sustainability. Aquac Eng 43:43–83.

54. Leonard N, Blancheton J, Guiraud J. 2000. Populations of heterotrophic bacteria in an experimental recirculating aquaculture system. Aquac Eng 22:22–109.

55. Schreier HJ, Mirzoyan N, Saito K. 2010. Microbial diversity of biological filters in recirculating aquaculture systems. Curr Opin Biotechnol 21:21–318.

56. Itoi S, Niki A, Sugita H. 2006. Changes in microbial communities associated with the conditioning of filter material in recirculating aquaculture systems of the pufferfish Takifugu rubripes. Aquaculture 256:256–287.

57. Burr GS, Wolters WR, Schrader KK, Summerfelt ST. 2012. Impact of depuration of earthy-musty off-flavors on fillet quality of Atlantic salmon, Salmo salar, cultured in a recirculating aquaculture system. Aquac Eng 50:50–28.

58. Blancheton JP, Attramadal KJK, Michaud L, d’Orbcastel ER, Vadstein O. 2013. Insight into bacterial population in aquaculture systems and its implication. Aquac Eng 53:53–30.

59. Kim D, Hofstaedter CE, Zhao C, Mattei L, Tanes C, Clarke E, Lauder A, Sherrill-Mix S, Chehoud C, Kelsen J, Conrad M, Collman RG, Baldassano R, Bushman FD, Bittinger K. 2017. Optimizing methods and dodging pitfalls in microbiome research. Microbiome 5:52.

60. Campbell JH, Foster CM, Vishnivetskaya T, Campbell AG, Yang ZK, Wymore A, Palumbo AV, Chesler EJ, Podar M. 2012. Host genetic and environmental effects on mouse intestinal microbiota. ISME J 6:6–2033.

61. Hildebrand F, Nguyen TLA, Brinkman B, Yunta RG, Cauwe B, Vandenabeele P, Liston A, Raes J. 2013. Inflammation-associated enterotypes, host genotype, cage and inter-individual effects drive gut microbiota variation in common laboratory mice. Genome Biol 14:R4.

62. Shephard KL. 1994. Functions for fish mucus. Rev Fish Biol Fish 4:4–401.

63. Ledy K, Giambérini L, Pihan JC. 2003. Mucous cell responses in gill and skin of brown trout Salmo trutta fario in acidic, aluminium-containing stream water. Dis Aquat Organ 56:235–240.

64. Vatsos IN, Kotzamanis Y, Henry M, Angelidis P, Alexis MN. 2010. Monitoring stress in fish by applying image analysis to their skin mucous cells. Eur J Histochem EJH 54.

65. Karlsen C, Ytteborg E, Timmerhaus G, Høst V, Handeland S, Jørgensen SM, Krasnov A. 2018. Atlantic salmon skin barrier functions gradually enhance after seawater transfer. Sci Rep 8.

66. Pittman K, Pittman A, Karlson S, Cieplinska T, Sourd P, Redmond K, Ravnøy B, Sweetman E. 2013. Body site matters: an evaluation and application of a novel histological methodology on the quantification of mucous cells in the skin of Atlantic salmon, Salmo salar L. J Fish Dis 36:36–115.

67. Tarnecki AM, Burgos FA, Ray CL, Arias CR. 2017. Fish Intestinal Microbiome: Diversity and Symbiosis Unraveled by Metagenomics. J Appl Microbiol n/a-n/a.

68. Dehler CE, Secombes CJ, Martin SAM. 2017. Environmental and physiological factors shape the gut microbiota of Atlantic salmon parr (Salmo salar L.). Aquaculture 467:467–149.

69. Wang C, Sun G, Li S, Li X, Liu Y. 2018. Intestinal microbiota of healthy and unhealthy Atlantic salmon Salmo salar L. in a recirculating aquaculture system. J Oceanol Limnol 36:36–414.

70. Bates JM, Mittge E, Kuhlman J, Baden KA, Cheesman SE, Guillemin K. 2006. Distinct signals from the microbiota promote different aspects of zebrafish gut differentiation. Developmental Biology 297: 374–386.

